# Controlled Release of Bone Morphogenetic Protein-2 Improves Motor Function After Traumatic Brain Injury in a Rat Model

**DOI:** 10.1101/2025.04.16.649206

**Authors:** Jakob M. Townsend, Jasmine Z. Deng, Scott Barbay, Brian T. Andrews, Randolph J. Nudo, Michael S. Detamore

**Affiliations:** C. Wayne McIlwraith Translational Medicine Institute, Colorado State University, Fort Collins, CO 80523; School of Biomedical Engineering, Colorado State University, Fort Collins, CO 80523; Bioengineering Graduate Program, University of Kansas, Lawrence, KS 66045; Department of Physical Medicine and Rehabilitation, University of Kansas Medical Center, Kansas City, KS 66103; Department of Otolaryngology, University of Iowa Medical Center, Iowa City, IA 52245; Department of Mechanical Engineering, Colorado State University, Fort Collins, CO 80523

**Author notes:** Corresponding Author: Jakob M. Townsend, PhD.

**Keywords:** BMP-2, Bone Regeneration, Extracellular Matrix, Hyaluronic Acid, Hydrogel, Traumatic Brain Injury

## Abstract

Severe traumatic brain injury (TBI) is a life-threatening condition characterized by internal brain swelling and commonly treated using a two-stage surgical approach. The interval between surgeries, generally spaced weeks to months, is associated with secondary neurologic complications from leaving the brain unprotected. Hydrogels may reshape severe TBI treatment by enabling a single-stage surgical intervention, capable of being implanted at the initial surgery, remaining flexible to accommodate brain swelling, and calibrated to regenerate bone after brain swelling has subsided. The current study evaluated the use of a pentenoate-modified hyaluronic acid (PHA) polymer with thiolated devitalized tendon (TDVT) for calvarial bone regeneration in a rat TBI model. Additionally, PHA-TDVT hydrogels encapsulating microspheres containing bone morphogenetic protein-2 (BMP-2) were investigated to enhance bone regeneration. All hydrogel precursor formulations exhibited sufficient yield stress for surgical placement. The addition of TDVT to the crosslinked hydrogels increased the average compressive modulus. *In vitro* cell studies revealed that the PHA-TDVT hydrogel with the highest concentration of BMP-2 microspheres (i.e., PHA-TDVT+µ100) significantly improved calcium deposition and osteogenic gene expression. Minimal *in vivo* bone regeneration was observed for all hydrogel groups; however, BMP-2 microsphere addition fortuitously reduced motor skill impairment and brain atrophy. The PHA-TDVT+µ100 group had 2.8 times greater reach index and 2.3 times lower brain atrophy values compared to the negative control (p<0.05). Overall, hydrogels with controlled release of BMP-2 may provide neuroprotective benefits in TBI treatment. Future studies should explore BMP-2 delivery strategies to enhance both bone and brain recovery in rat TBI studies.

**Graphical Abstract:** 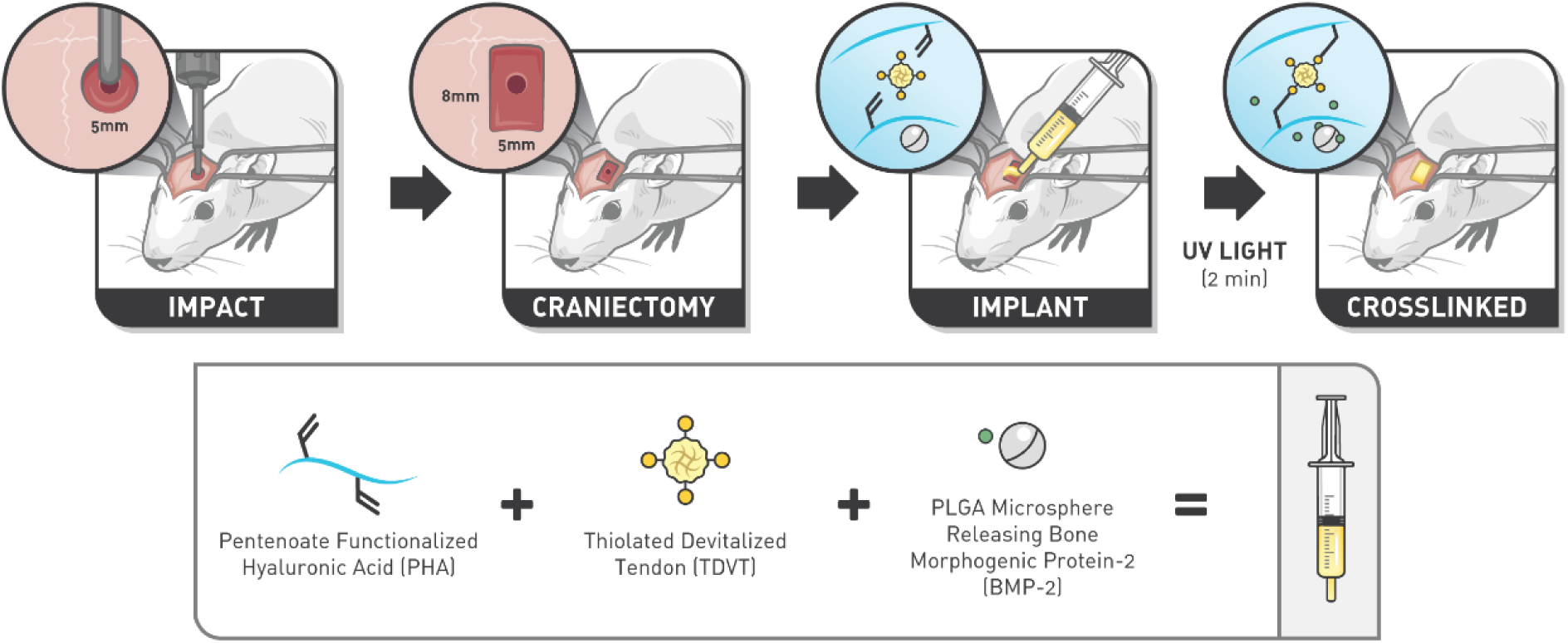

**Statement of Significance:** Severe traumatic brain injury (TBI) is a life-threatening condition characterized by internal brain swelling and is commonly treated using a two-stage surgical approach. Complications associated with the two-stage treatment paradigm include secondary neurologic impairment, termed syndrome of the trephined (SOT). SOT is often reversible once the second surgery is performed, whereas a single-stage TBI treatment paradigm may avoid the occurrence of SOT altogether. Utilizing hydrogels comprised of pentenoate-modified hyaluronic acid and thiolated devitalized tendon encapsulating microspheres containing bone morphogenetic protein-2 (BMP-2), the current study demonstrated improvements in motor skill function and reductions in brain atrophy in a rat TBI model. The introduction of hydrogels with controlled release of BMP-2 as a neuroprotective strategy for TBI application offers a promising approach for single-stage TBI treatment.

## Introduction

Severe traumatic brain injury (TBI) is a serious and potentially life-threatening condition characterized by internal brain swelling.^1, 2^ Decompressive craniectomy (DC) is a lifesaving procedure commonly used to treat closed skull TBI. DC treatment reduces intracranial pressure by removing a large portion of bone, permitting the brain to swell beyond the confines of the calvarial vault.^3^ After brain swelling has subsided, cranioplasty is performed to rebuild the calvarial vault. The timing between surgeries may be artificially lengthened due to clinical scheduling or clinician preference, with reports ranging from weeks to several months.^4^ An inherent issue with the typical two-stage surgical intervention is the prevalence of secondary neurologic conditions associated with leaving the brain unprotected.^5^ Termed syndrome of the trephined (SOT) or sinking skin flap syndrome, patients may suffer from headache, dizziness, mood swings, and/or fine motor dexterity issues during the time between surgeries.^6^ Although the cause of SOT remains unclear, the condition is often reversible once cranioplasty is performed.^7, 8^ An ideal surgical approach for TBI treatment would eliminate the second surgery altogether, avoiding neurological conditions associated with an unprotected brain, and reducing patient recovery time.^9^

To successfully eliminate the need for a second surgery, both the brain injury and the DC bone defect must be addressed during the initial—and therefore, only—surgery. One approach to effective TBI treatment involves using a biomaterial that supports cranial bone regeneration over time while allowing for brain swelling and promoting recovery. To that end, hydrogels offer a unique biomaterial solution for TBI treatment, as their inherent flexibility provides distinct advantages. Additionally, there is a need for a biomaterial that can easily flow into the defect, conform to patient-specific contours, and set rapidly to minimize procedural time constraints ensuring retention during healing.^10^

Current research on TBI treatment after DC has primarily focused on reducing inflammation and promoting brain healing.^11–16^ Few studies have explored hydrogels as a strategy to reduce the number of surgeries during TBI treatment.^17, 18^ Current clinical products and methods are unsuitable for single-stage TBI treatment as they do not remain pliable after implantation. Hydrogels are an attractive choice for single-stage TBI treatment due to their tunable post-crosslinking stiffness and inherent flexibility, offering advantages over existing clinical products. Hydrogels designed for bone regeneration over a specified time interval are well-suited for TBI treatment, as they remain flexible during brain swelling and gradually transition to bone. A single-stage hydrogel approach also avoids leaving the brain unprotected, potentially reducing neurologic complications associated with the current two-stage surgical paradigm.^19–21^

The biomaterial design in the current study was motivated by the need to overcome limitations of current biomaterials for single-stage TBI treatment. The objective of the current study was to evaluate a pentenoate-functionalized hyaluronic acid (PHA) polymer with thiolated devitalized tendon (TDVT) tissue particles capable of crosslinking to form an interconnected hydrogel for cranial bone restoration in a TBI rat model. Additionally, bone morphogenetic protein (BMP)-2 was loaded into poly(lactic-co-glycolic acid) microspheres that were embedded in the hydrogel to enhance bone formation. The photocrosslinking hydrogels presented in the current study build upon our previously published work as a next-generation biomaterial.^22–24^ The advantage of the hydrogel platform lies in the surface modification of devitalized tendon particles with thiol groups, enabling TDVT to act as the crosslinker and facilitating rapid covalent bonding with the PHA polymer.^24^ Thiol-ene photopolymerization chemistry is a preferred alternative to methacrylate-based systems in the operating room due to its rapid crosslinking of only 1-2 min.^25–27^

The rationale for incorporation of TDVT tissue particles in the current study was an attempt to recapitulate part of the endochondral ossification process by providing an intermediate raw material to accelerate bone formation.^28^ Devitalized tendon is rich in collagen I, and our previous work demonstrated that PHA-DVT effectively promoted bone regeneration compared to demineralized bone matrix.^22^ In the current study, PHA hydrogels with TDVT concentrations of 5, 10, and 15% were selected for *in vitro* characterization of the constructs. To assess the paste-like consistency for surgical application, rheological analysis of the hydrogel precursor solution (pre-crosslinking) was conducted to determine yield stress and recovery after disruption. Success criteria for yield stress (>100 Pa) and recovery (>80%) were selected to ensure ease of material placement, as we discussed in a review of hydrogel precursor rheology for surgical placement.^29^ To evaluate material retention after surgical placement, mechanical characterization of the crosslinked hydrogel was conducted to measure swelling ratio, absorption, and stiffness. Success criteria for hydrogel stiffness (>100 kPa) ensured adequate material retention at the defect site. To evaluate the bioactivity of the hydrogel for cranial bone regeneration, *in vitro* cell studies were performed to determine the leading TDVT concentration and assess the synergistic benefit of controlled BMP-2 delivery.

Finally, the leading TDVT formulation was evaluated in a rat TBI model with recovery assessed using micro-computed tomography and histology for bone regeneration and brain atrophy. Additionally, motor skill recovery was evaluated using a skilled reach test. Control groups included: (1) DC alone (craniectomy without TBI), (2) DC+TBI (no hydrogel treatment), (3) TBI closed skull (DC+TBI with immediate cranioplasty), and (4) PHA alone (no TDVT incorporation, material control). We hypothesized that the TDVT hydrogel with controlled release of BMP-2 would promote greater bone regeneration and motor recovery compared to control groups.

## Methods

### Thiolation of devitalized tendon particles

Porcine Achilles tendon tissue (6 months, Male, Berkshire) was obtained from Oklahoma State University College of Veterinary Medicine from unrelated studies. Tendon tissue was cut into 5 mm cubes using a scalpel, coarse-ground using a cryogenic tissue grinder (BioSpec Products, Bartlesville, OK), then frozen at −20⁰C and lyophilized. Tendon tissue was then cryoground using a freezer-mill (SPEX 6775, SamplePrep, Metuchen, NJ) to produce devitalized tendon (DVT) and stored at −20⁰C for later use. DVT particles were thiolated using our previously published method.^23^ Briefly, DVT was weighed dry and suspended at a concentration of 5 mg/mL in dissolving buffer comprised of 50 mM sodium phosphate (Cat# 342483, Sigma-Aldrich, St. Louis, MO) and 1 mM ethylenediaminetetraacetic acid (EDTA, Cat# EDS, Sigma-Aldrich) at pH 7.5. N-succinimidyl S-acetylthioacetate (Cat# 573100, Sigma-Aldrich) was then dissolved in dimethylformamide (DMF, Cat# 227056, Sigma-Aldrich) at a concentration of 100 mM, then 2 mL of the DMF solution was added to the DVT solution to initiate the thiolation reaction. The thiolation reaction proceeded for 1 hour under constant mixing at room temperature. The reacted DVT solution was then filtered through Whatman 1 (Cat# WHA1001090, Sigma-Aldrich) filter paper, where the reacted DVT particles were retained on the filter paper surface. Reacted DVT particles were collected and resuspended in dissolving buffer at a concentration of 5 mg/mL assuming no loss of DVT particles. Acetylated-thiols on the reacted DVT particles were then deprotected using deprotecting solution to create the TDVT. The deprotecting solution was comprised of 500 mM hydroxylamine hydrochloride (Cat# 159417, Sigma-Aldrich), 50 mM sodium phosphate, and 25 mM EDTA at pH 7.5. 20 mL of deprotecting solution was added to the dissolving buffer for every gram of DVT. The deprotecting solution was allowed to react with the reacted DVT for 2 hours at room temperature under constant mixing. Afterward, the resulting TDVT was dialyzed (MWCO: 6-8 kDa, Cat# 08-670F, Thermo Fisher Scientific, Waltham, MA) against deionized (DI) water for 48 hours, performing exchanges every 12 hours. The TDVT solution was frozen, lyophilized, and stored dry at −20°C for later use. DVT thiolation was confirmed using a modified Ellman’s assay. Approximately 5 mg of TDVT tissue particles were weighed dry and suspended in 600 µL of phosphate buffered saline (PBS, Cat# P3813, Sigma-Aldrich). Ellman’s reagent (5-5’-dithiobis(2-nitrobenzoic acid), Cat# D218200, Sigma-Aldrich) was dissolved at a concentration of 1 mg/mL in PBS, and then 100 µL of the dissolved Ellman’s reagent was added to the suspended tissue particles. The reaction was allowed to proceed for 5 mins while vortexing every 30 s to ensure complete mixing. Afterward, the reaction solution was centrifuged at 12,000 × g, and then 100 µL of the resulting supernatant was transferred to a 96 well plate and the absorbance was read at 412 nm. A standard curve of L-cysteine (Cat# W326305, Sigma-Aldrich) was prepared by serially diluting a 1 mM stock to determine unknown sample concentrations.

### Pentenoate-modified hyaluronic acid synthesis

Hyaluronic acid (HA) was modified by conjugating pentenoate groups using our previously published method.^24^ Briefly, HA (Mw = 1.64 MDa, Cat# HA15M-5, Lifecore Biomedical, Chaska MN) was dissolved in DI water at a concentration of 0.5% (w/v) for 2 h. After complete dissolution, DMF was added dropwise to a final ratio of 3:2 (DI:DMF). Next, the catalyst 4-(dimethylamino)pyridine was added to the solution to a final concentration of 6 mM. The pH was then adjusted to 9 using 1 M NaOH (Cat# S5881, Sigma-Aldrich). To initiate the reaction, pentenoic anhydride (Cat# 471801, Sigma-Aldrich) was added dropwise to reach a 5 M excess relative to HA while maintaining the pH between 8 and 9 using 1 M NaOH for 2 hours. Once the pH maintained a constant value, indicating completion of the reaction, the pentenoate-modified HA (PHA) was precipitated in 5 volumes of acetone and then centrifuged at 6000 × g. The supernatant was then decanted and the PHA pellets were dissolved in DI water at a concentration of 1% (w/v). The dissolved PHA was then transferred to dialysis tubing (MWCO: 6-8 kDa) and dialyzed against DI water for 96 h, performing DI water exchanges every 24 h. After the dialysis phase, the PHA solution was frozen, lyophilized, and stored at −20°C. Addition of pentenoate groups to the HA backbone was confirmed using NMR (Varian VNMRS-500 MHz NMR spectrometer, Varian, Palo Alto, CA) as previously described.^30^

### Hydrogel preparation and crosslinking

Hydrogels were prepared using our previously described method.^31^ Briefly, PHA and TDVT were weighed dry and combined. PHA-only hydrogels contained 4% (w/v) PHA without addition of TDVT, and composite hydrogels with 4% PHA and 5, 10, or 15% TDVT were referred to as PHA-TDVT hydrogels. For studies evaluating the PHA-TDVT hydrogel with BMP-2 microspheres, the formulation was comprised of 4% PHA and 15% TDVT with 1 (PHA-TDVT+µ1), 10 (PHA-TDVT+µ10), or 100 (PHA-TDVT+µ100) mg of BMP-2 microspheres per mL of hydrogel. Dry materials were sterilized using ethylene oxide gas (AN74i, Anderson Anprolene, Haw River, NC). The sterilized dry components were resuspended in sterile PBS containing 2.3 mM lithium phenyl-2,4,6-trimethylbenzoylphosphinate (LAP, Cat# 900889, Sigma-Aldrich) and 5 mM dithiothreitol (DTT, Cat# D0632, Sigma-Aldrich). The PBS-LAP-DTT solution was sterile-filtered using a 0.22 µm syringe filter prior to use. Hydrogel precursor solutions were allowed to fully dissolve for 2 h prior to crosslinking. For *in vitro* characterization and cell studies, hydrogels were fabricated using a previously published protocol.^32^ Briefly, hydrogel precursor mixtures were syringed into 1 mm thick silicon molds sandwiched between glass microscope slides. Hydrogel precursors were then crosslinked using 365 nm UV- light (Cat# EA-160, 1280 µW/cm^2^, Spectro-UV, Farmingdale, NY) on both sides for 1 min each. Hydrogels were then punched into cylindrical discs using a 6 mm biopsy punch. For *in vivo* studies, hydrogels were spread into the defect using a spatula and crosslinked *in situ* for 2 min using the same UV-light used in the *in vitro* studies.

### Rheological characterization

The yield stress (n = 3) and percent recovery of G’ (n = 3) for the hydrogel precursor solutions prior to crosslinking were characterized using a DHR-2 controlled stress rheometer (TA Instruments, New Castle, DE). A single batch for each group was created and evaluated three times total for each method. Rheological measurements were performed using a 20 mm diameter flat stainless steel plate fixture and Peltier plate cover at 37°C and a gap distance of 500 μm. For yield stress testing of the hydrogel precursor, an oscillatory shear stress sweep at a frequency of 1 Hz was used. The stress corresponding to the crossover point of the storage (G’) and loss (G”) modulus was defined as the yield stress.^30^ To determine the percent recovery of G’ for the hydrogel precursor solutions, samples were subjected to a constant shear stress of 10 Pa for 5 min to establish the baseline, then sheared at a stress above the material’s yield stress for 30 s, then returned to a shear stress of 10 Pa. Precursor solutions were sheared at a constant frequency of 1 Hz during each phase of the percent recovery of G’ experiment. The storage modulus baseline was defined as the average storage modulus value during low shear and the recovered storage modulus was defined as the storage modulus 5 s after the end of the high stress phase.

### Hydrogel absorption and swelling ratio

PHA-TDVT hydrogels post-crosslinking were evaluated to determine the impact of incorporating thiolated tissue particles on the absorption and swelling characteristics. The PHA hydrogel served as the negative control, and experimental TDVT concentrations of 5, 10, and 15% were selected for evaluation. Hydrogel absorption (n = 6) and swelling (n = 6) ratios were determined as follows. The initial hydrogel mass (m_i_) was recorded directly after crosslinking in silicon molds. Hydrogels were then swollen to equilibrium in PBS at 37°C for 24 h and the swollen mass (m_s_) was recorded. Afterward, hydrogels were frozen at −20°C, lyophilized for 48 h, then the dry hydrogel mass (m_d_) was recorded. Hydrogel absorption was calculated as the ratio of the swollen mass to the initial hydrogel mass (Eq. 1) and the swelling ratio was calculated as the ratio of the swollen mass to dry mass (Eq. 2).

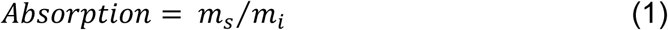

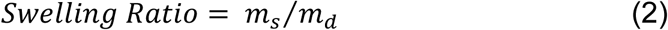

The compressive moduli of the hydrogels (n = 6) after crosslinking were determined using a DHR-2 controlled stress rheometer (TA Instruments). Crosslinked hydrogels were prepared using a previously published method and explained in the hydrogel preparation and crosslinking section above.^33^ After fabrication, hydrogels were swelled in PBS for 24 h. Once swelling was complete, the swollen hydrogel diameter was measured using a micrometer and the height was measured at room temperature under dry conditions using the DHR-2 rheometer at a tare load of 0.01 N. Hydrogels were compressed at a constant rate of 5 μm/s (~0.25% strain/s) until 20% strain. The compressive modulus was calculated from the linear portion of the stress-strain curve between 5 and 15% strain.

### Microsphere fabrication and release study

Microspheres were produced using a microencapsulator (Encapsulator B-390, Buchi, New Castle, DE) equipped with a concentric nozzle consisting of a 120 µm inner nozzle (i.e., core) and a 300 µm outer nozzle (i.e., shell). Immediately prior to encapsulation, 1 g of poly(lactic-co-glycolic acid) (PLGA) was dissolved in 9 mL of dichloromethane (DCM, Cat# 320269, Sigma-Aldrich), and 1 mg of recombinant human BMP-2 (INFUSE™, Medtronic, Dublin, Ireland) was dissolved in 1 mL of sterile phosphate buffered saline. The two solutions were combined and sonicated at an amplitude of 30 µm for 30 s, creating a fine emulsion with a BMP-2 loading of 1 mg/g PLGA. The shell stream and hardening bath were composed of 1% polyvinyl alcohol (PVA, 25 kDa, 88% hydrolyzed, Cat# 02975, Warrington, PA) dissolved in deionized water. The polymer-protein emulsion and PVA solution were loaded into separate syringes, placed in syringe pumps (NE-1000, New Era Pump Systems Inc., Farmingdale, NY), and carefully attached to the core and shell inlets of the concentric nozzle device.

The core solution was pumped at a rate of 1.5 mL/min and the shell solution at a rate of 6 mL/min to form a stable stream. The electrode (1500 V) and vibrational generator (1500 Hz) were then activated to begin microsphere formation. microspheres were collected in 500 mL of the hardening bath solution under constant agitation at 200 rpm. After the droplet collection step, microspheres were allowed to set for 3 hours before collecting, washing with 5 L of deionized water, freezing, lyophilizing, and then storing at −20⁰C for later use.

Release of BMP-2 from PLGA microspheres was assessed over a 42-day period using a BMP-2 enzyme-linked immunosorbent assay (ELISA) kit (Cat# EHBMP2, Thermo Fisher Scientific) following the manufacturer’s protocol (n = 3). A single batch of BMP-2 microspheres was produced and tested in triplicate. Approximately 20 mg of BMP-2 microspheres were weighed dry and transferred to 1.5 mL centrifuge tubes with 1 mL of 0.1% bovine serum albumin (BSA, Cat# A9418, Sigma-Aldrich) dissolved in PBS. Microcentrifuge tubes were placed on a tube rotator and incubated at 37⁰C for 42 days. At time points of 1, 3, 7, 10, 14, 21, 28, 35, and 42 days, the supernatant was extracted and replaced with fresh PBS+BSA solution. Extracted supernatants were stored in sterile microcentrifuge tubes at −20⁰C for later quantification.

### In vitro cell culture

Human bone marrow-derived mesenchymal stem cells (hBMSCs, 19 years old, Male, RoosterBio Inc, Frederick, MD) expanded in RoosterNourish™ (RoosterBio Inc) supplemented with 1% penicillin/streptomycin (P/S, Cat# 15140-122, Thermo Fisher Scientific) were passaged twice before use. After >80% cell confluence, hBMSCs were extracted and transferred to minimum essential medium-α (α-MEM, Cat# 12561072, Thermo Fisher Scientific) supplemented with 10% certified fetal bovine serum (FBS, Cat# 16000044, Thermo Fisher Scientific), 1% P/S, 10 mM β-glycerophosphate (β-gly, Cat# G9422, Sigma-Aldrich), and 250 µM ascorbic acid-2-phosphate (A2P, Cat# 49752, Sigma-Aldrich). α-MEM supplemented with FBS, P/S, β-gly, and A2P was used as the negative control medium. For the positive control medium, the base negative control medium with the addition of 100 ng/mL human recombinant BMP-2 (Cat# 355-BM, R&D Systems, Minneapolis, MN) was used for osteogenic differentiation. Experimental hydrogels were prepared as described in the hydrogel preparation and crosslinking section above. Crosslinked hydrogel discs were placed in the wells of tissue culture treated 96 well plates and swelled for 24 h in 200 μL of PBS at 37°C. The PBS was aspirated and then 10,000 cells/well (66,666 cells/mL) were added in the negative or positive control medium. For the cell-only control groups, 10,000 cells/well (66,666 cells/mL) were added into the wells of tissue culture treated 96 well plates without hydrogel. After 10 days of cell culture (n = 12 wells/group), cell culture medium was aspirated from the wells and stored in microcentrifuge tubes at −20°C for osteocalcin quantification. Samples intended for reverse transcription quantitative PCR (RT-qPCR) analysis were incubated with 200 μL of lysis buffer (Cat# R1060-50, Zymo Research, Irvine, CA) for 10 mins and then stored at −80°C for later use. Samples intended for calcium deposition quantification were incubated in 1M hydrochloric acid for 24 hours at 37°C then stored at −20°C for later use.

### In vitro assays

Calcium content was assayed using the QuantiChrom™ calcium assay kit (Cat# DICA-500, BioAssay Systems, Hayward, CA) according to the manufacturer’s protocol (n = 6). Calcium standards were prepared in 1M hydrochloric acid to match the experimental samples. Osteocalcin content in cell culture media was assayed using an ELISA (Cat# LS-F3640, Lifespan Biosciences Inc., Newark, CA) according to the manufacturer’s protocol (n = 6). RT-qPCR was performed using a CFX Opus 96 Real-Time PCR System (Bio-Rad, Hercules, CA). RNA was isolated using the *Quick*-RNA Miniprep Kit (Cat# R1051, Zymo Research, Irvine, CA) and transcribed to cDNA using the iScript™ Reverse Transcription Supermix (Cat# 1708840, Bio-Rad). PCR amplification was performed using the iQ™ Multiplex Powermix (Cat# 1725849, Bio-Rad) and the following probes (Bio-Rad): GAPDH (Cat# qHsaCEP0041396), Collagen I (COLI, Cat# qHsaCEP0050510), Osteocalcin (OCN, Cat# qHsaCEP0041159), and RUNX2 (Cat# qHsaCEP0051329). Six samples from each group (n = 6) were tested in duplicate. Relative levels of gene expression were calculated using the **ΔΔ**C_T_ method as previously described.^33^ Briefly, hBMSCs cultured in negative control medium were designated as the calibrator group and GAPDH was used as the endogenous control.

### Rat Skilled Reach Training and Testing

Animal experiments were approved by the Institutional Animal Care and Use Committee of the University of Kansas Medical Center. A total of 58 male Long-Evans rats (10-12 weeks old) were purchased from Envigo (Summerset, NJ) and trained on the skilled reach task prior to surgery and subsequent skilled reach testing. Prior to the experiment, 12 of the 58 rats were used to determine parameters for a cortical impact that could produce a long-term impairment of fine motor skills over a two-month recovery period. During the experiment there was a loss of 7 rats from the remaining 46 utilized for the study: 6 rats did not meet pre-operative training criteria and there was one post-operative fatality. A total of 39 rats completed pre-operative training and were used for experimental testing. Rats were housed in pairs upon arrival within a climate-controlled vivarium maintained at 20-22⁰C with a 12 hr /12 hr light-dark cycle and allowed a 48-hour acclimation period prior to handling. Rats were handled for 1-2 weeks prior to being individually housed at the beginning of behavioral training. Water was provided ad libitum throughout the experiment and food was scheduled to be available after behavioral sessions to provide sufficient motivation for training and assessment. Each rat’s normal intake of rodent biscuits (Harlan Teklad Rodent Diet 8604) was calculated at >3% of their body weight, which was measured 3 times per week. Supplemental food was provided if a rat’s weight dropped by >15%.

Rats were assessed on the skilled reach test before and after the surgical procedure to evaluate motor performance, as we have previously published.^34^ The skilled reaching task required rats to reach from inside of a behavioral testing apparatus to grasp and retrieve a small food pellet (45mg Rodent Dustless Precision Pellets; Bio-Serv Inc., Frenchtown, NJ) from a shelf attached to an outside wall. The apparatus consisted of a 12” by 12” Plexiglas box with two 15 mm wide openings placed on the left and right side of the front wall providing access to a Plexiglas shelf placed at a distance of 30 mm from the openings. Openings were cut 30 mm from each edge, allowing the rats to only use the preferred forelimb to retrieve a pellet. An automated door prevented reaching before a food pellet was in place. The door was controlled by a linear actuator (Actuonix, Vitoria, BC) activated by an infrared beam break sensor placed at the rear of the testing apparatus. A second infrared beam break sensor at the front of the box activated the doors to close once the forelimb retracted. Each rat received 60 trials per timepoint, with one food pellet delivered per trial. Training and testing sessions typically required 15-20 minutes per rat. A webcam attached to the side wall recorded each reach. A reach was considered successful when the rat extended its forelimb through the opening, grasped the food pellet from the shelf, and retrieved the pellet to their mouth without dropping it.^35^ Rats were trained 5 days per week over a 3-week period, where inclusion criteria required >60% proficiency. Reach values were collected every other day for 3 consecutive sessions and averaged to establish pre-injury baseline values. Rats unable to achieve >60% reaching proficiency were excluded from the study. Post-surgical reach performance was assessed once per week for 8 weeks. A performance index was used to normalize each rat’s post-injury performance to their average pre-injury baseline. A performance index of 100% indicated post-injury performance was equal to or had returned to average baseline performance. An index value <100% indicated post-injury motor impairment, while an index value >100% indicated improved performance from baseline. An arcsine transformation was used to normalize data for statistical analysis.^7^

### Rat surgical procedure and traumatic brain injury

After establishing pre-injury baseline performance values for skilled reach testing, rats were randomly assigned an identification number for group placement and reassigned a different number after the surgical procedure to mask group assignment. Thus, investigators conducting the behavioral testing and subsequent offline skill assessment from recorded videos were blinded to group assignment. Surgical procedures were performed under aseptic conditions, followed by daily post-operative pain management for three days post-surgery (buprenorphine 0.05 mg/kg subcutaneous. and acetaminophen 20 mg/kg by mouth) and daily medical monitoring for the first week. The average weight of the rats at the time of surgery was 384 ± 14 g. Each rat was anesthetized in an induction chamber with 3% isoflurane and then moved to a breathing cone delivering 1-2% isoflurane for the duration of the procedure. Anesthesia depth was monitored every 15 minutes by observing whisking, pinch reflex, or blinking. Body temperature was monitored throughout the procedure using a rectal probe and maintained with a Kent Scientific Far infrared warming pad (~36⁰C to 37⁰C). Ophthalmic ointment (Puralube Vet Ointment, Dechra Veterinary, Overland Park, KS) was applied to the eyes to mitigate desiccation. The head was shaved, cleaned with alcohol and positioned in a stereotaxic frame, secured in the prone position. Bupivacaine (0.5 mL subcutaneously) was administered along the intended surgical area for analgesic effects, followed by another cleaning of the scalp with alcohol and iodine. A 0.15 mL injection of penicillin was provided subcutaneously (45K IU G benzathine and G procaine). The scalp was incised at the midline using a scalpel and retracted to expose the skull. The periosteum tissue was removed, and the temporal muscle retracted to expose the calvarial bone.

To assess the efficacy and sustainability of treatment effects over an extended period, 12 rats were initially used to determine cortical impact parameters required to create a chronic fine motor impairment (i.e., diameter, depth, velocity, and duration of the impact). A cortical impact resulting in a chronic impairment of skilled e forelimb use without impairing gross motor function was achieved. Preserving gross motor movement was necessary for participation in behavioral testing during the acute and chronic recovery phases. For the overall TBI procedure, an initial craniectomy was performed to expose the brain and enable TBI induction. Immediately after TBI, the craniectomy was expanded to mimic DC. All rats received a 5 mm diameter craniectomy over the hemisphere opposite to the preferred forelimb (i.e., the forelimb most often used during the skilled reach task) to expose the caudal forelimb area (CFA; primary motor cortex homolog in rats).^7^ Stereotaxic coordinates for the initial and expanded craniectomies were marked with a sterile surgical pen. The initial 5 mm diameter craniectomy was created using a 5 mm outer diameter dental trephine (Fine Science Tools) centered over the M1 forelimb motor cortex (i.e., CFA; 1 mm rostral to Bregma and 3 mm lateral to midline).^36^ A TBI was induced using a controlled cortical impact (CCI) technique, centered within the 5 mm craniectomy. The impact was delivered using an electromagnetic CCI device (Impact One, Leica Biosystems, Buffalo Grove, IL) mounted on a stereotaxic frame to reproduce the position and direction of the impact with high precision. A 4 mm diameter stainless-steel impact rod with a beveled edge was used to prevent the dura from tearing. The impact was delivered to a depth of 3 mm from the surface of the dura at 4.5 m/s with an impact time of 200 msec. Immediately after CCI, DC groups received an expanded 5 by 8 mm rectangular craniectomy centered around the initial craniectomy using Littauer bone cutters to score the calvaria and rongeurs to remove the bone.

For cranial repair, either a cranioplasty using standard acrylic (Jet™ Denture Repair Acrylic Resin Professional Package, Cat# 1323-C, Lang Dental Manufacturing Co Inc., Wheeling IL) or 50 µL of hydrogel formulation was syringed into the bone defect immediately after DC for rats receiving cranial repair or left unrepaired. Control groups included: DC (i.e., DC without TBI, n = 7), DC+TBI (i.e., no hydrogel treatment, n = 6), TBI closed (i.e., DC+TBI with immediate cranioplasty using standard acrylic, n = 6), and PHA (i.e., no TDVT incorporation, material control, n = 7) were employed to accurately assess experimental hydrogel impact. Experimental hydrogel groups included PHA-TDVT (i.e., 4% PHA and 15% TDVT, n = 7) and PHA-TDVT+µ100 (i.e., 4% PHA, 15% TDVT, and 100 mg BMP-2 microspheres/mL hydrogel, n = 6). A 365 nm ultraviolet light (Cat# EA-160, 1280 µW/cm^2^, Spectro-UV) was used to crosslink hydrogel formulations once applied to the cranial opening and cured for 2 minutes prior to closing. The scalp was then sutured using 4-0 silk and treated with bupivacaine, eutectic mixture of local anesthetics (EMLA) cream, and triple antibiotics (Neosporin). After 10 weeks post-CCI, rats were humanely euthanized following AVMA guidelines.

### Micro-computed tomography (µCT)

Micro-computed tomography (µCT) imaging of explanted rat calvarial defects and brain tissue was performed after 10 weeks of recovery. For the rat calvarial defects, a Quantum GX2 imaging system (PerkinElmer, Waltham, MA) with a 50 kV X-ray source at 160 μA produced a voxel size of 36 μm (n = 6 or 7). Avizo software (FEI Company, Hillsboro, OR) was used to quantify bone volume within the defect. Regenerated bone was quantified within a rectangular volume of interest (VOI) of 8 mm in length and 5 mm in width, with the volume centered over the calvarial defect. Bone samples from age-, breed-, and sex-matched animals from unrelated studies were used to define a minimum global threshold value to enable quantification of new bone within the VOI. Regenerated bone within the defect was delineated blue and quantified (mm^3^).

For quantification of the CCI injury within brain tissue, a contrast enhanced μCT imaging method was adapted to better visualize soft tissue.^37, 38^ Briefly, brain tissue was submerged in 50 mL of a graded series of ethanol (i.e., 30, 50, 70, 90%) for 2 hours at room temperature prior to incubating the tissue in contrast solution (i.e., 90% methanol and 1% ioversol) for 24 hours. Brain tissue was then rehydrated using 70 and 30% ethanol for 2 hours each before submerging the tissue in deionized water and imaging. For μCT imaging of brain tissue, a Quantum FX (PerkinElmer) imaging system with a 90 kV X-ray source at 200 μA produced a voxel size of 40 μm (n = 6 or 7). A 10 mm VOI centered over the defect and spanning both hemispheres of the brain was utilized for calculating the percent atrophy volume. The injured hemisphere was demarcated red, and the uninjured hemisphere was demarcated blue. Percent brain atrophy was defined as the percentage volume difference between the injured and uninjured hemispheres.

### Histological analysis

Rat calvarial bone samples for histological analysis were harvested and stored in 10% phosphate-buffered formalin for 48 h (n = 6 or 7). Embedding, sectioning, and staining were performed using standard methods. Briefly, explanted calvarial bone samples were decalcified using Newcomer Supply Decalcifying Solution (Cat# NC2101886, Fisher Scientific, Hampton, NH) at room temperature for 2 weeks. Decalcified bone was embedded in paraffin and sectioned in the sagittal plane at a thickness of 4 µm and mounted on glass microscope slides. Tissue slides were deparaffinized in standard gradients of xylene and ethanol to water prior to staining. Hematoxylin and eosin (H&E) staining was performed using the SelecTech H&E staining system (Leica Biosystems, Wetzlar, Germany) on a Leica ST5020 multistainer (Leica Biosystems) following the manufacturer’s protocol.

Explanted brain samples were removed from the skull and submerged in 10% phosphate buffered formalin for 48 h (n = 6 or 7). Embedding, sectioning, and staining were performed using standard methods. Briefly, explanted brains were transferred to a histology matrix (Cat# 15065, Ted Pella INC, Redding, CA) and a 2 mm thick coronal section was taken. The anterior extent of the section was approximately midway between the anterior and posterior limits of the brain defect (i.e., midline of the CCI impact). High profile microtome blades were used to maintain a symmetrical cut across the tissue. Sections were placed anterior side down in tissue cassettes and transferred to individual specimen containers containing 50 mL of 70% ethanol. Brain tissue was embedded in paraffin and sectioned in the coronal plane to a thickness of 15 µm and affixed to glass microscope slides. Tissue slides were deparaffinized in standard gradients of xylene and ethanol to water prior to staining. For cresyl violet staining, slides were incubated in 0.1% cresyl violet (Cat# C5042, Sigma-Aldrich) for 5 min, briefly rinsed in DI water for 10 s, then dipped in 1% acetic acid. Differentiation in the stained tissue was achieved by submerging the slides in 95% ethanol repeatedly until the desired staining was achieved.

### Statistical methods

Statistical analyses were conducted using GraphPad Prism (GraphPad Software Inc., La Jolla, CA). A one-way analysis of variance (ANOVA) was used to analyze the pre- and post-crosslinking hydrogel mechanical evaluation, *in vitro* studies involving microspheres, and *in vivo* studies. A two-way ANOVA was used to analyze the *in vitro* study assessing the impact of medium containing BMP-2. Hydrogel precursor rheology and microsphere release testing had n = 3 and all other *in vitro* testing had n = 6. *In vivo* testing had n = 6 or 7 samples per group. Results are reported as the mean ± standard deviation.

## Results

### Rheological analysis of hydrogel precursor

All hydrogel formulations tested exhibited a detectable yield stress (Fig. 1A). The yield stress of the PHA-TDVT 15% formulation (1070 ± 1.9 Pa) was 2.5, 2.6, and 1.5 times greater compared to PHA, PHA-TDVT 5%, and PHA-TDVT 10%, respectively (p<0.05). The yield stress of the PHA-TDVT 10% formulation (730 ± 150 Pa) was 1.7 and 1.8 times greater compared to PHA and PHA-TDVT 5%, respectively (p<0.05). No significant differences in yield stress were observed among the other groups. All hydrogel precursors exhibited >80% recovery of the storage modulus (G’) after shearing (Fig. 1B). No significant differences in G’ recovery were observed among the groups, which ranged from 88 to 98% after 5 s of recovery time. The highest average recovery time was observed in the PHA-TDVT 15% formulation (98%).

**Figure 1.**
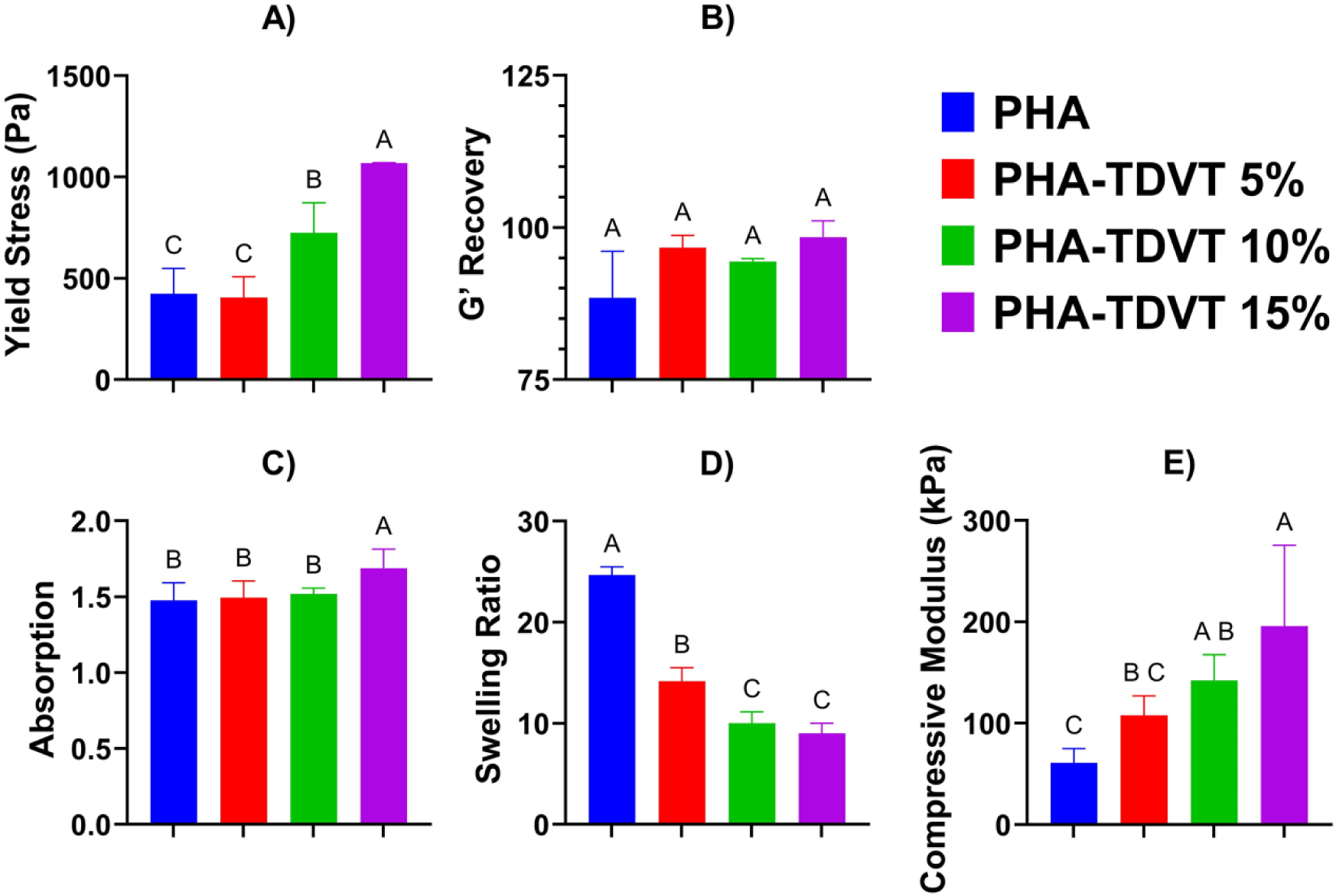
Hydrogel precursor rheology and post-crosslinking mechanical characterization. A) Yield stress of the hydrogel precursor determined by the crossover point of the storage (G’) and loss (G”) modulus (n = 3). B) Precursor recovery determined by the restoration of the storage modulus (G’) after shearing (n = 3). C) Crosslinked hydrogel absorption ratio demonstrating the change in mass after fabrication (n = 6). D) Swelling ratio of crosslinked hydrogels demonstrating the ratio of water to polymer mass indicative of the crosslinking density (n = 6). E) Compressive modulus of crosslinked hydrogels determined by the slope of the stress-strain curve between 5 and 15% (n = 6). Note that the PHA-TDVT 15% group exhibited the greatest average yield stress, G’ recovery, and compressive modulus. PHA = pentenoate functionalized hyaluronic acid. TDVT = thiolated devitalized tendon. Letters indicate significant differences from other groups with different letters (A>B>C, p<0.05).

### Mechanical characterization of crosslinked hydrogels

All hydrogel formulations exhibited increased water content at equilibrium (Fig. 1C). The hydrogel absorption ratio of the PHA-TDVT 15% formulation (1.7 ± 0.13) was 1.1 times greater compared to PHA, PHA-TDVT 5%, and PHA-TDVT 10% (p<0.05). No significant differences in absorption were observed among the other groups. Increasing TDVT particle content in the PHA hydrogel matrix decreased the average hydrogel swelling ratio (Fig. 1D). The hydrogel swelling ratio for the PHA group (25 ± 0.8) was 1.7, 2.5, and 2.7 times greater compared to PHA-TDVT 5%, PHA-TDVT 10%, and PHA-TDVT 15%, respectively (p<0.05). Additionally, the swelling ratio of the PHA-TDVT 5% group (14 ± 1.4) was 1.4 and 1.6 times greater compared to PHA-TDVT 10% and PHA-TDVT 15%, respectively (p<0.05). No significant differences in swelling were observed among the other groups.

All PHA-TDVT formulations demonstrated an adequate compressive modulus (i.e., >100 kPa) for application to TBI treatment (Fig. 1E). The compressive modulus of the PHA-TDVT 15% group (200 ± 80 kPa) was 3.2 and 1.8 times greater compared to PHA and PHA-TDVT 5%, respectively (p<0.05). In addition, the compressive modulus of the PHA-TDVT 10% group (140 ± 25 kPa) was 2.3 times greater compared to PHA (p<0.05). No significant differences in compressive modulus were observed among the other groups.

### TDVT concentration and BMP-2 synergy in vitro

Increasing TDVT concentration with and without soluble BMP-2 addition demonstrated greater average calcium content (Fig. 2A). Calcium content ranged from 3.2 to 6.8 µg for groups cultured without soluble BMP-2 and 3.7 to 7.4 µg for groups cultured with soluble BMP-2. The greatest average calcium content for groups cultured without BMP-2 was observed in the PHA-TDVT 15% group (6.8 ± 0.4 µg), which was 2.1, 1.7, 1.3, and 1.2 times greater than calcium contents of the TCP, PHA, PHA-TDVT 5%, and PHA-TDVT 10% groups, respectively (p<0.05). The greatest average calcium content for groups cultured with BMP-2 (i.e., +BMP-2) was observed in the PHA-TDVT 15% group (7.4 ± 0.7 µg), which was 2.0, 1.7, 1.3, and 1.2 times greater compared to TCP, PHA, PHA-TDVT 5%, and PHA-TDVT 10%, respectively (p<0.05). Although average calcium content was greater for each group cultured with BMP-2, no significant differences were observed for individual groups between medium conditions.

**Figure 2.**
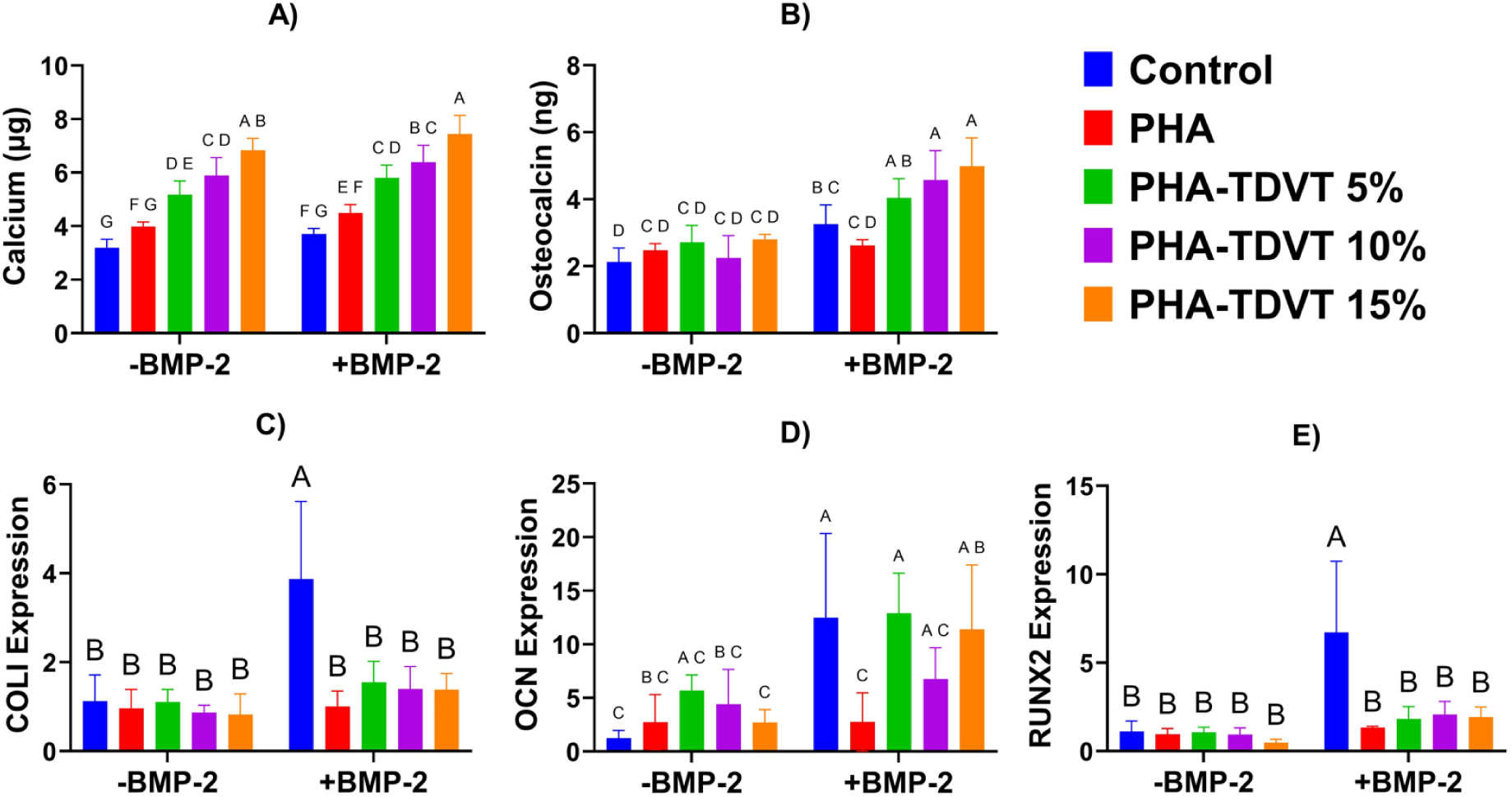
*In vitro* response of human bone marrow-derived mesenchymal stem cells (hBMSCs) seeded on PHA hydrogels with increasing TDVT content (n = 6). Seeded cells were cultured in medium with and without BMP-2 to explore synergy between the PHA-TDVT hydrogel and osteogenic signaling. A) Intracellular and deposited calcium content after 10 days of culture. B) Cell culture media osteocalcin protein content secreted by hBMSCs in response to increasing hydrogel TDVT content and media condition. C-E) Gene expression of hBMSCs for COLI, OCN, and RUNX2. Note a significantly higher value for calcium content and osteocalcin secretion for the PHA-TDVT 15% group compared to the base PHA hydrogel and cells cultured on tissue culture plastic in the presence of BMP-2 (p<0.05). BMP-2 = bone morphogenetic protein-2. PHA = pentenoate functionalized hyaluronic acid. Control = tissue culture plastic. TDVT = thiolated devitalized tendon. Letters indicate significant differences from other groups with different letters (A>B>C>D>E>F>G, p<0.05).

A synergistic improvement in average osteocalcin content was observed for the PHA-TDVT groups cultured with BMP-2 compared to without (Fig. 2B). Osteocalcin content ranged from 2.1 to 2.6 ng for groups cultured without BMP-2 and 2.5 to 4.1 ng for groups cultured with BMP-2. For the medium condition without BMP-2, no differences were observed among groups. For the medium condition with BMP-2, the greatest average osteocalcin content was observed in the PHA-TDVT 15% group (4.1 ± 0.7 ng). Osteocalcin content for the PHA-TDVT 15% (+BMP-2) group (4.1 ± 0.7 ng) was 1.9, 1.8, 1.6, 1.8, 1.6, 1.2, and 1.6 times greater compared to TCP (−BMP-2), PHA (−BMP-2), PHA-TDVT 5% (−BMP-2), PHA-TDVT 10% (−BMP-2), PHA-TDVT 15% (−BMP-2), TCP (+BMP-2), and PHA (+BMP-2), respectively (p<0.05). For the PHA-TDVT 10% (+BMP-2) group (4.0 ± 0.7 ng), osteocalcin content was 1.9, 1.8, 1.5, 1.8, 1.5, 1.2, and 1.6 times greater compared to TCP (−BMP-2), PHA (−BMP-2), PHA-TDVT 5% (−BMP-2), PHA-TDVT 10% (−BMP-2), PHA-TDVT 15% (−BMP-2), TCP (+BMP-2), and PHA (+BMP-2), respectively (p<0.05). Finally, osteocalcin content for the PHA-TDVT 5% (+BMP-2) group (3.6 ± 0.5 ng) was 1.7, 1.6, 1.4, 1.6, 1.4 and 1.5 times greater compared to TCP (−BMP-2), PHA (−BMP-2), PHA-TDVT 5% (−BMP-2), PHA-TDVT 10% (−BMP-2), PHA-TDVT 15% (−BMP-2), and PHA (+BMP-2), respectively (p<0.05). No differences in osteocalcin content were observed among PHA-TDVT groups.

Gene expression of hBMSCs seeded on PHA-TDVT hydrogels confirmed expression of osteocalcin (Fig. 2C-E). No differences among groups for COLI (Fig. 2C) and RUNX2 (Fig. 2E) gene expression were observed among groups, with the exception of the TCP (+BMP-2) group. For COLI expression, the TCP (+BMP-2) group (3.9 ± 1.8) was 3.4, 4.0, 3.5, 4.5, 4.7, 3.9, 2.5, 2.8, and 2.8 times greater compared to TCP (−BMP-2), PHA (−BMP-2), PHA-TDVT 5% (−BMP-2), PHA-TDVT 10% (−BMP-2), PHA-TDVT 15% (−BMP-2), PHA (+BMP-2), PHA-TDVT 5% (+BMP-2), PHA-TDVT 10% (+BMP-2), and PHA-TDVT 15% (+BMP-2), respectively (p<0.05). For RUNX2 expression, the TCP (+BMP-2) group (6.7 ± 4.0) was 6.0, 7.0, 6.4, 7.2, 13.8, 5.0, 3.7, 3.2, and 3.5 times greater compared to TCP (−BMP-2), PHA (−BMP-2), PHA-TDVT 5% (−BMP-2), PHA-TDVT 10% (−BMP-2), PHA-TDVT 15% (−BMP-2), PHA (+BMP-2), PHA-TDVT 5% (+BMP-2), PHA-TDVT 10% (+BMP-2), and PHA-TDVT 15% (+BMP-2), respectively (p<0.05). Elevated average osteocalcin gene expression values for TCP, PHA-TDVT 5%, and PHA-TDVT 15% cultured with BMP-2 were observed (Fig. 2D). Osteocalcin gene expression values for the TCP (+BMP-2) group (13 ± 7.9) were 10.1, 4.5, 2.8, 4.6, and 4.5 times greater compared to TCP (−BMP-2), PHA (−BMP-2), PHA-TDVT 10% (−BMP-2), PHA-TDVT 15% (−BMP-2), and PHA (+BMP-2), respectively (p<0.05). Osteocalcin gene expression values for the PHA-TDVT 5% (+BMP-2) group (13 ± 3.7) were 10.5, 4.7, 2.9, 4.8, and 4.7 times greater compared to TCP (−BMP-2), PHA (−BMP-2), PHA-TDVT 10% (−BMP-2), PHA-TDVT 15% (−BMP-2), and PHA (+BMP-2), respectively (p<0.05). Finally, osteocalcin gene expression values for the PHA-TDVT 15% (+BMP-2) group (11 ± 6.4) were 9.2, 4.2, 4.2, and 4.1 times greater compared to TCP (−BMP-2), PHA (−BMP-2), PHA-TDVT 15% (−BMP-2), and PHA (+BMP-2), respectively (p<0.05). No other significant differences in osteocalcin gene expression values were observed among the other groups.

### PHA-TDVT hydrogel with BMP-2 microspheres in vitro

Cumulative release of BMP-2 from PLGA microspheres demonstrated an initial burst release after day 1 followed by a secondary burst release after 21 days (Fig. 3A). At timepoints of 1, 3, 7, 14, 21, 28, 35, and 42 days the percentage of BMP-2 payload delivery was 1.7, 9.8, 2.2, 0.7, 1.1, 63.1, 20.4, and 1.0 %, respectively. Synergistic improvement in calcium deposition (Fig. 3B) and osteogenic gene expression (Fig. 3C-E) was observed for the PHA-TDVT 15% hydrogel with 100 µg/mL of BMP-2 microspheres (i.e., PHA-TDVT+µ100). Calcium deposition for the PHA-TDVT+µ100 group (2.8 ± 0.9 µg) was 2.6, 2.5, 1.9, and 1.6 times greater compared to PHA-TDVT, PHA-TDVT+BMP-2, PHA-TDVT+µ10, and PHA-TDVT+µ50, respectively (p<0.05). No significant differences in calcium content were observed among the other groups. COLI gene expression for the PHA-TDVT+µ100 group (4.1 ± 1.9) was 4.1, 2.1, 2.7, and 2.1 times greater compared to PHA-TDVT, PHA-TDVT+BMP-2, PHA-TDVT+µ10, and PHA-TDVT+µ50, respectively (p<0.05). No significant differences in COLI gene expression were observed among the other groups. OCN gene expression for the PHA-TDVT+µ100 group (47 ± 2.2) was 46, 9.8, 23, and 2.4 times greater compared to PHA-TDVT, PHA-TDVT+BMP-2, PHA-TDVT+µ10, and PHA-TDVT+µ50, respectively (p<0.05). Additionally, OCN gene expression for the PHA-TDVT+µ50 group (20 ± 2.0) was 19, 4, and 9.5 times greater compared to PHA-TDVT, PHA-TDVT+BMP-2, PHA-TDVT+µ10, respectively (p<0.05). Finally, OCN gene expression for the PHA-TDVT+BMP-2 group (4.8 ± 1.9) was 4.7 and 2.3 times greater compared to PHA-TDVT and PHA-TDVT+µ10, respectively (p<0.05). No significant differences in OCN gene expression were observed among the other groups. RUNX2 gene expression for the PHA-TDVT+µ100 group (3.3 ± 0.9) was 3.3, 2.2, 2.2, and 1.4 times greater compared to PHA-TDVT, PHA-TDVT+BMP-2, PHA-TDVT+µ10, and PHA-TDVT+µ50, respectively (p<0.05). Additionally, RUNX2 gene expression for the PHA-TDVT+µ50 group (2.4 ± 0.9) was 2.3 times greater compared to PHA-TDVT (p<0.05). No significant differences in RUNX2 gene expression were observed among the other groups.

**Figure 3.**
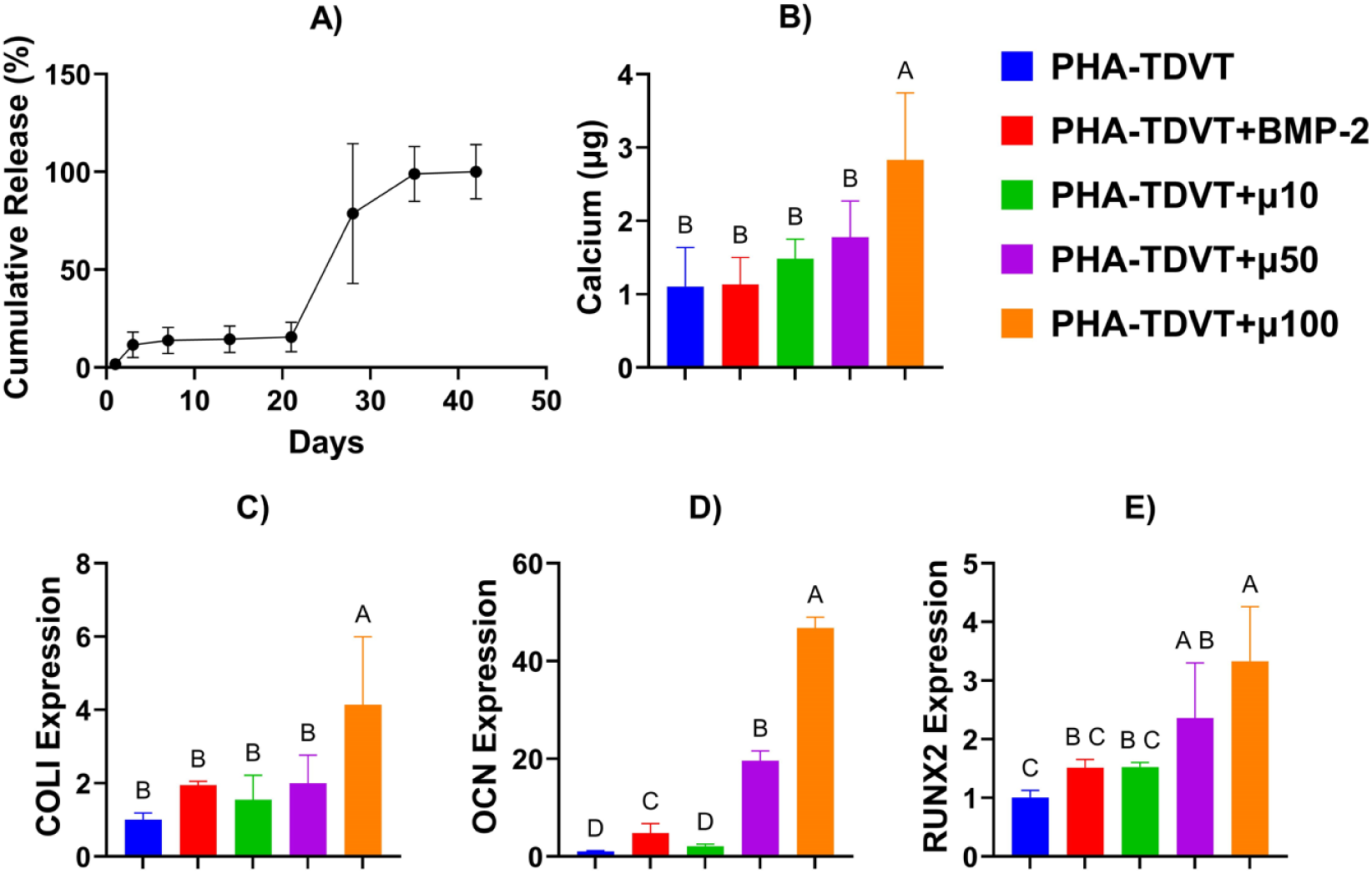
BMP-2 release profile and response of human bone marrow mesenchymal stem cells (hBMSCs) seeded onto PHA-TDVT hydrogels with 10, 50, or 100 mg/mL of BMP-2 microspheres. A) Release profile of BMP-2 microspheres over 42 days (n = 6). Note an initial burst phase followed by a secondary release after 21 days. B) Calcium content of PHA-TDVT hydrogels after 14 days of culture with seeded hBMSCs (n = 8). C-E) Gene expression of hBMSCs for COLI, OCN, and RUNX2 after 14 days of culture (n = 6). Note that the PHA-TDVT+µ100 (100 mg/mL BMP-2 microspheres) group exhibited the greatest average calcium content and highest gene expression for COLI, OCN, and RUNX2. PHA = pentenoate functionalized hyaluronic acid. TDVT = thiolated devitalized tendon. BMP-2 = bone morphogenetic protein-2. PHA-TDVT+BMP-2 = PHA-TDVT cultured with soluble BMP-2 (100 ng/mL). µ10 = 10 mg BMP-2 microspheres/mL hydrogel. Letters indicate significant differences from other groups with different letters (A>B>C>D, p<0.05).

### Bone microcomputed tomography

New bone formation at the periphery of the defect site was observed in all groups (Fig. 4A). Additionally, bone island formation was observed in all groups. Quantification of bone regeneration ranged from averages of 9.1 to 16.9 mm^3^ (Fig. 4B). The PHA group (16.9 ± 3.7 mm^3^) had 1.8 times greater new bone volume compared to PHA-TDVT+µ100 (p<0.05). No significant differences in bone regeneration were observed among the other groups.

**Figure 4.**
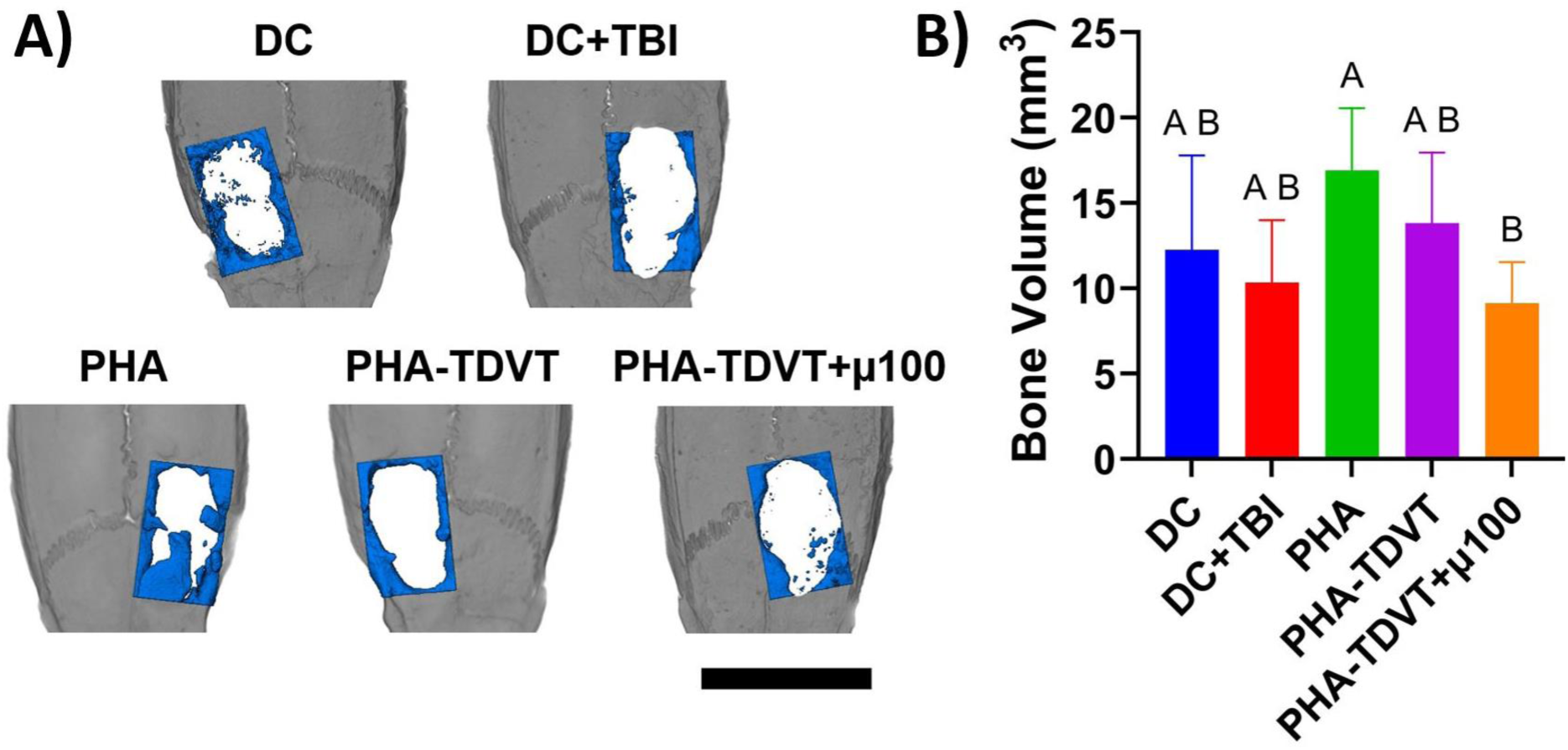
Microcomputed tomography reconstructions and bone volume evaluation using Avizo software after 10 weeks of recovery (n = 6 or 7). An initial square defect measuring 8 by 5 mm (L × W) was created on the calvarial surface and 50 µL of hydrogel was implanted into the empty defect. A) Representative microcomputed tomography reconstructions with regenerated bone colored in blue. Scale bar = 10 mm. B) Quantified bone volume within the region of interest. Note that the lowest average bone regeneration was observed in the PHA-TDVT+µ100 group. DC = decompressive craniectomy without traumatic brain injury (TBI). DC+TBI = decompressive craniectomy with TBI. PHA = pentenoate functionalized hyaluronic acid. TDVT = thiolated devitalized tendon. µ100 = 100 mg BMP-2 microspheres/mL hydrogel. Letters indicate significant differences from other groups with different letters (A>B, p<0.05).

### Bone histological analysis

New bone formation was observed in all experimental groups spanning into the defect as determined by H&E staining (Fig. 5). Soft tissue formation was observed in the defect site of all groups where new bone was absent. New bone formation observed in all groups was absent of necrotic bone formation. No inflammatory cells or inflammatory reactive tissue were observed in the experimental groups.

**Figure 5.**
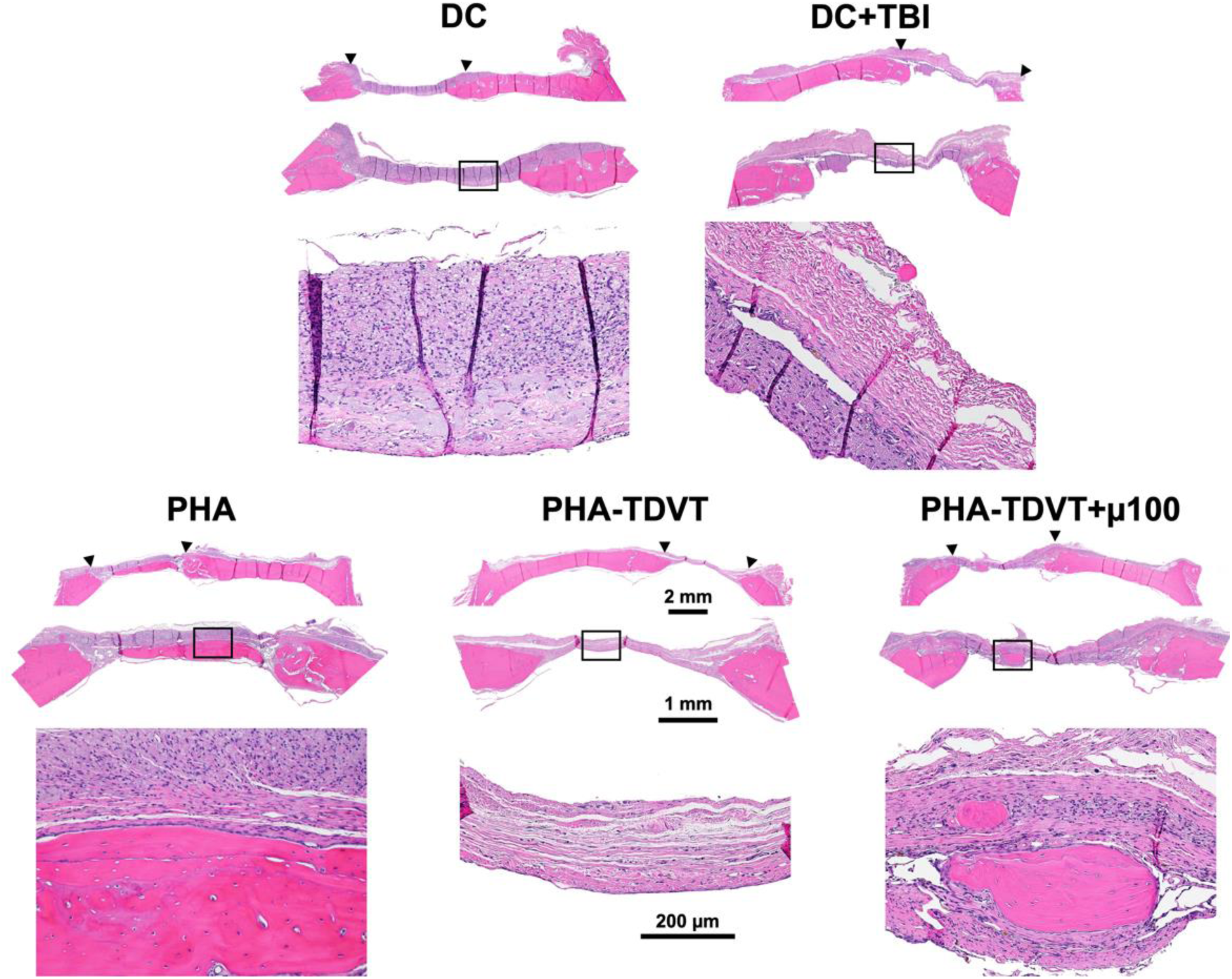
Hematoxylin and eosin (H&E) histological staining of rectangular (8×5 mm) rat cranial defects 10 weeks after implantation (n = 6 or 7). Sections were taken in the coronal plane with the dural side of the calvarium positioned toward the bottom of each image. Under each group name, the full cranial section appears on top with an expanded view of the defect denoted by arrows shown in the middle image. A magnified view of the defect space within the black rectangle is shown in the bottom image. Note peripheral bone growth and absence of necrotic bone formation in all groups. DC = decompressive craniectomy without traumatic brain injury (TBI). DC+TBI = decompressive craniectomy with TBI. PHA = pentenoate functionalized hyaluronic acid. TDVT = thiolated devitalized tendon. µ100 = 100 mg BMP-2 microspheres/mL hydrogel.

### Skilled reach testing analysis

Skilled reach testing for all experimental TBI groups demonstrated an initial decline in motor function after TBI to varying extents (Fig. 6). The skilled reach index for the DC group (i.e., no TBI) demonstrated minimal impairment in motor function after surgery, as expected (Fig. 6A). All experimental TBI groups demonstrated improvement in reach testing over the course of the 8-week period, with the exception of the PHA group. The aggregate skilled reach index over the course of 8 weeks demonstrated less impairment in the PHA-TDVT+µ100 group compared to the other experimental TBI groups (Fig. 6B). The aggregate reach index for the DC group (88 ± 2.7%) was 4.4, 2.6, 4.7, 4.5, and 1.6 times greater compared to DC+TBI, TBI Closed, PHA, PHA-TDVT, and PHA-TDVT+µ100, respectively (p<0.05). For the PHA-TDVT+µ100 group (57 ± 9.2%), the aggregate reach index was 2.8, 1.7, 3.0, and 2.9 times greater compared to DC+TBI, TBI Closed, PHA, and PHA-TDVT, respectively (p<0.05). Finally, the aggregate reach index for the TBI Closed group (34 ± 11%) was 1.7, 1.8, and 1.7 times greater compared to DC+TBI, PHA, and PHA-TDVT, respectively (p<0.05). No significant differences in aggregate reach index were observed among the other groups.

**Figure 6.**
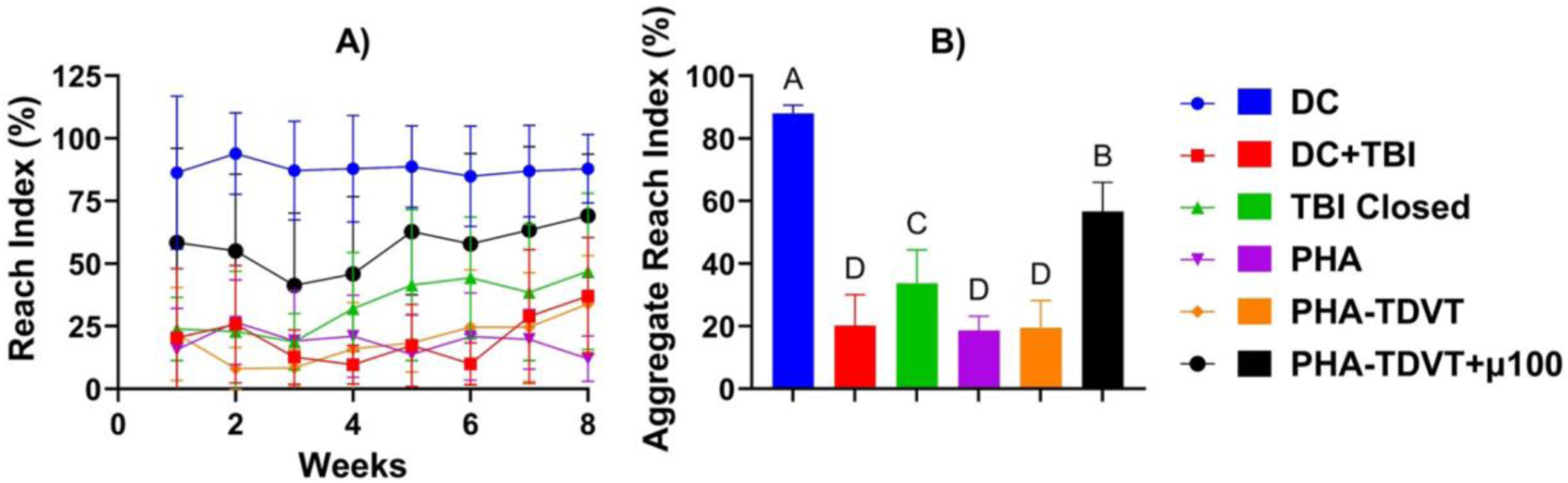
Motor skill recovery evaluated over 8 weeks of recovery (n = 6 or 7). A) Weekly reach index testing using the skilled reach test method. Note an initial average decline in reach index for all groups, with the greatest degree of recovery observed in the PHA-TDVT+µ100 group. B) Aggregate skilled reach testing over the course of 8-weeks. Note a significantly greater value for the aggregate reach index for the PHA-TDVT+µ100 group compared to the other experimental TBI groups (p<0.05). DC = decompressive craniectomy without traumatic brain injury (TBI). DC+TBI = decompressive craniectomy with TBI. PHA = pentenoate functionalized hyaluronic acid. TDVT = thiolated devitalized tendon. µ100 = 100 mg BMP-2 microspheres/mL hydrogel. Letters indicate significant difference from other groups with different letters (A>B>C>D, p<0.05).

### Brain microcomputed tomography

Brain atrophy at the CCI impact site was observed in all experimental groups that received the TBI procedure, as determined by microcomputed tomography (Fig. 7). Brain atrophy and degree of tissue volume loss varied within the VOI (Fig. 7A) among the experimental conditions (Fig. 7B). The DC group (0.5 ± 2.0 %) group exhibited 21, 16, 15, and 14 times lower brain atrophy values compared to DC+TBI, TBI Closed, PHA, and PHA-TDVT groups, respectively (p<0.05). Additionally, the brain atrophy percentage for the PHA-TDVT+µ100 group (4.5 ± 1.2 %) was 2.3 times lower compared to the DC+TBI group (p<0.05). No significant differences in brain atrophy were observed among other groups.

**Figure 7.**
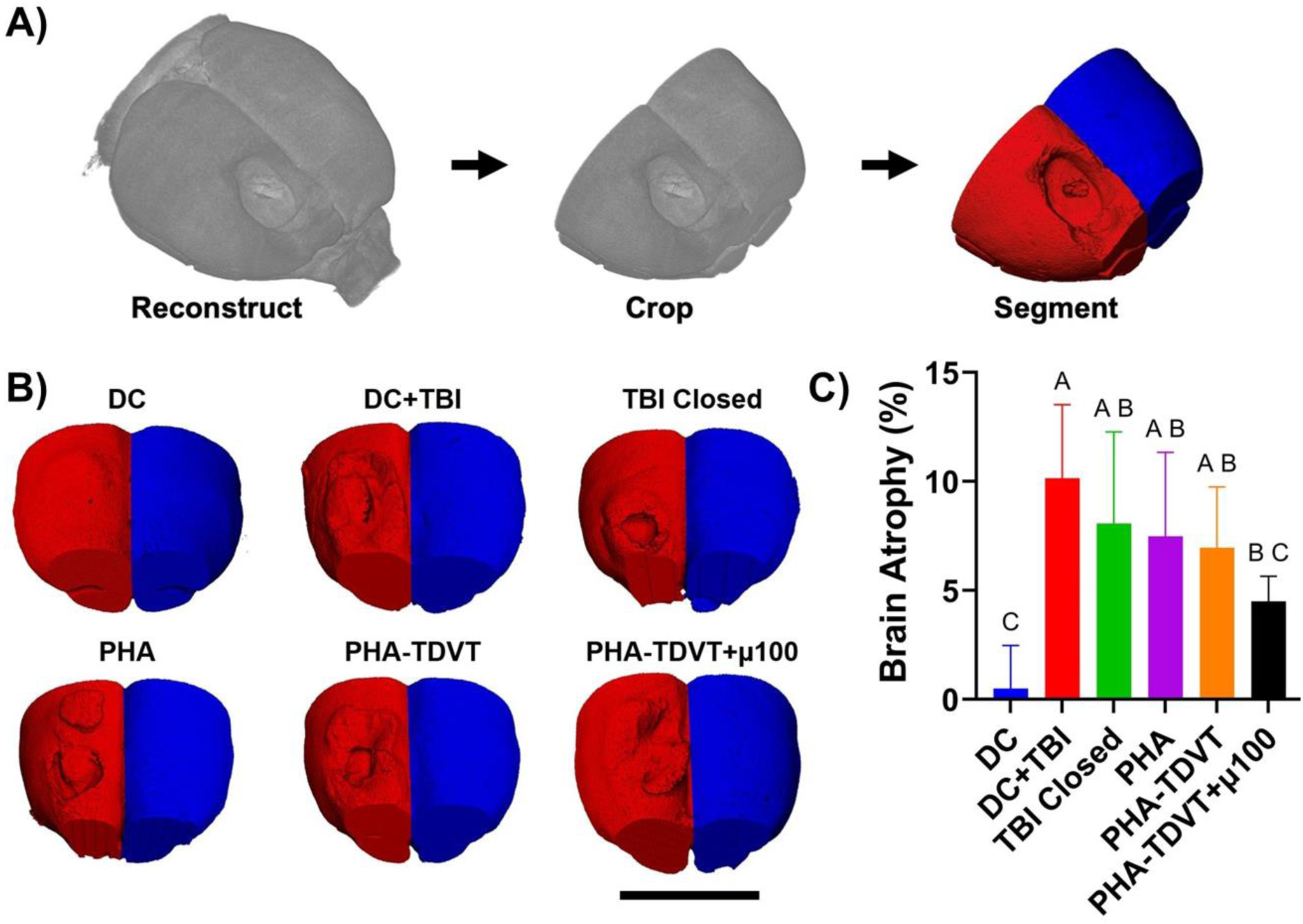
Brain microcomputed tomography reconstructions and quantification of atrophy volume (n = 6 or 7). A) Demonstration of the microcomputed tomography evaluation where scans were reconstructed, cropped within a 10 mm region of interest, then segmented into the uninjured or injured hemisphere. B) Microcomputed tomography reconstructions of experimental groups, note the presence of brain atrophy after injury. Scale bar = 10 mm. C) Quantification of brain atrophy by normalizing the injured hemisphere to the uninjured hemisphere. Note that the PHA-TDVT+µ100 group exhibited the lowest average brain atrophy value compared to the other experimental groups. DC = decompressive craniectomy without traumatic brain injury (TBI). DC+TBI = decompressive craniectomy with TBI. TBI Closed = decompressive craniectomy and TBI with immediate cranioplasty using dental acrylic. PHA = pentenoate functionalized hyaluronic acid. TDVT = thiolated devitalized tendon. µ100 = 100 mg BMP-2 microspheres/mL hydrogel. Letters indicate significant differences from other groups with different letters (A>B>C, p<0.05).

### Brain histological analysis

Brain atrophy and tissue loss were observed in all experimental groups as determined by cresyl violet staining (Fig. 8). Additionally, brain swelling was observed in the DC group that did not receive the TBI procedure. Expansion of the ventricle space adjacent to the CCI impact site was observed to varying degrees in all experimental TBI groups. Brain atrophy and tissue loss localized to the neocortex was observed in the experimental groups with lesser extent observed in the PHA-TDVT+µ100 group.

**Figure 8.**
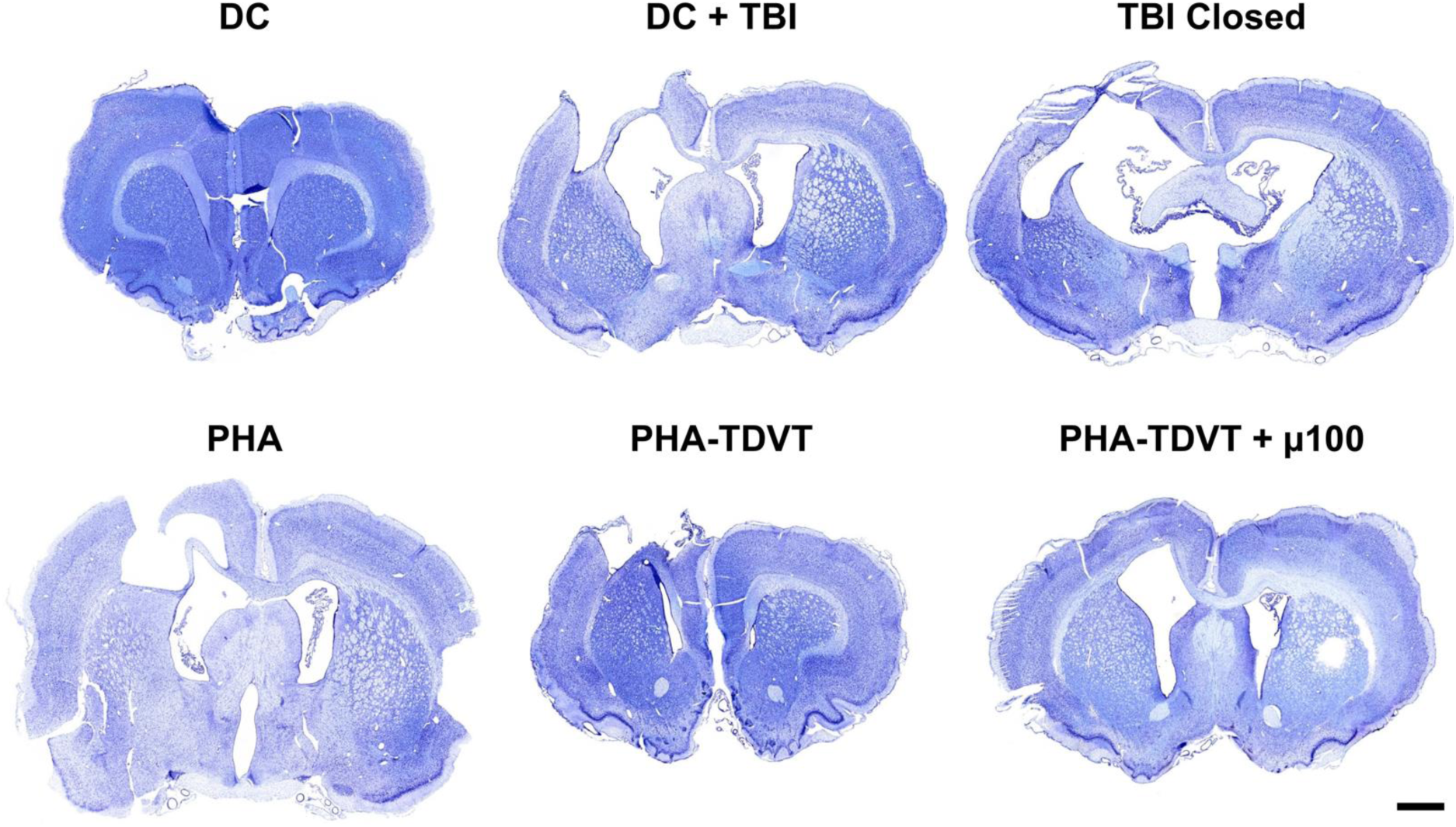
Cresyl violet histological staining of coronal brain sections. Coronal sections were taken from the midline of the visible brain lesion across the length of the tissue (n = 6 or 7). Note the presence of brain atrophy observed by the reduction in hemisphere area at the site of injury in TBI groups. Scale bar = 10 mm. DC = decompressive craniectomy without traumatic brain injury (TBI). DC+TBI = decompressive craniectomy with TBI. TBI Closed = decompressive craniectomy and TBI with immediate cranioplasty using dental acrylic. PHA = pentenoate functionalized hyaluronic acid. TDVT = thiolated devitalized tendon. µ100 = 100 mg BMP-2 microspheres/mL hydrogel.

## Discussion

The current study is the first to utilize a hydrogel construct for single-stage treatment of TBI following decompressive craniectomy in a rat model. The hydrogels introduced in the current study were composed of PHA polymer and TDVT particles that crosslink to form an interconnected matrix. The selection of devitalized tendon extracellular matrix (ECM) in the current study was based on our previous work demonstrating superior bone regeneration compared to demineralized bone matrix.^22^ The PHA-TDVT hydrogels exhibited desirable paste-like handling characteristics, defect shape-fitting properties, and rapid crosslinking times for surgical placement. Material characteristics, particularly mechanical performance in both fluid (pre-crosslinking) and solid (post-crosslinking) states, were identified as critical for future TBI treatment and clinical translation.^29^ To evaluate the feasibility of material placement, hydrogel precursor solutions were characterized using yield stress and recovery time experiments. Additionally, post-crosslinking characterization, including hydrogel compressive modulus, absorption, and swelling ratio, confirmed the material’s retention capability. All hydrogel precursor solutions exhibited sufficient yield stress (>100 Pa) and recovery time (>80% recovery G’), supporting their feasibility for material placement. For context, the yield stress of mayonnaise and Play-Doh is 200 and 3000 Pa, respectively.^22^ Following crosslinking, increasing TDVT content reduced the average swelling ratio and improved the average compressive modulus; however, these differences were not statistically significant across all formulations. The observed increase in compressive modulus with higher particle content was speculated to result from interactions between the PHA polymer and the thiolated tendon particles. Overall, the characterization results support the potential application of PHA-TDVT hydrogels for calvarial defect repair and TBI treatment.

A key anticipated observation in the current study was the synergy between PHA-TDVT hydrogel formulations and soluble BMP-2 for enhancing osteogenic gene expression and biochemical content *in vitro*. However, surgical implantation of materials with soluble BMP-2 has limitations due to the lack of a controlled and confined *in vitro* environment, where we have previously observed minimal bone regeneration in rat calvarial bone defects.^10^ To address concerns regarding *in vivo* BMP-2 delivery, we further explored controlled release of BMP-2 from PLGA microspheres to isolate and prolong BMP-2 to the defect site. Prior to further testing, the PHA-TDVT 15% hydrogel formulation was selected based on superior average calcium and osteocalcin content *in vitro*. As anticipated, PHA-TDVT hydrogels with BMP-2 microsphere delivery *in vitro* demonstrated improved biochemical content and osteogenic gene expression with increasing microsphere content. Moreover, the PHA-TDVT hydrogel with the greatest BMP-2 microsphere content significantly outperformed the positive control of the PHA-TDVT hydrogel with soluble BMP-2 (100 ng/mL) *in vitro*. Due to these observations, the PHA-TDVT+µ100 group was selected for *in vivo* implantation.

Our hypothesis that the PHA-TDVT hydrogel with controlled BMP-2 release would promote greater bone regeneration compared to the control groups was inconclusive. Overall, minimal *in vivo* performance differences were observed among the experimental groups in terms of regenerated bone volume and H&E staining. In contrast to our previous studies evaluating PHA-TDVT in 8 mm rat calvarial bone defects,^23^ the volume of regenerated bone in the current study was significantly lower. This discrepancy may be attributed to differences in defect geometry (i.e., circular versus rectangular) or the presence of a secondary TBI injury. Although all experimental groups exhibited low levels of bone regeneration, an unexpected finding was that the lowest average bone regeneration occurred in the PHA-TDVT group containing BMP-2 microspheres (i.e., PHA-TDVT+µ100). Possible explanations include a delay in BMP-2 release missing the optima therapeutic window, an insufficient BMP-2 dose, release of acidic byproducts from PLGA degradation impeding healing, or inadequate BMP-2 localization at the defect site. Despite the lack of substantial bone formation in the current study, BMP-2 administration yielded promising secondary outcomes, particularly in motor skill recovery and reduction of brain atrophy.

In contrast to bone regeneration outcomes, the PHA-TDVT+µ100 group demonstrated significant improvements in motor skill recovery (as assessed by skilled reach testing) and reduction in brain atrophy volume (measured via microcomputed tomography) compared to the control group of DC+TBI. Among the experimental TBI groups, the PHA-TDVT+µ100 group exhibited less impairment in average reach index one week after TBI. Although average brain atrophy was lower in the PHA-TDVT+µ100 group compared to other experimental groups, the difference was only significantly different when compared to the control DC+TBI group. Cresyl violet staining further supported the microcomputed tomography findings, revealing predominately intact neocortical tissue in the PHA-TDVT+µ100 group. The results of the current study suggest that BMP-2 may serve a dual role in both bone and brain recovery following TBI. Dual delivery of BMP-2, targeting both bone regeneration via conjugation to the material and brain recovery via controlled release, may help overcome limitations observed in the current study.

Other groups have investigated the use of BMP’s for neurological recovery across a wide array of applications.^39–42^ Zanella *et al*.^43^ evaluated the effects of BMP-2 applied to an absorbable collagen sponge implanted adjacent to or wrapped around the sciatic nerve to assess pain-associated behaviors. The aforementioned study unexpectedly observed neuroprotective effects of BMP-2, although the effect diminished over time. Compared to the current study, the prolonged neuroprotective impact observed may be attributed to the controlled release by microsphere delivery. In another study, Saglam *et al*.^44^ investigated the effects of BMP-2 on neurodegeneration in neuronal cells and primary neurons *in vitro*, observing that BMP-2 exerted neurotrophic effects through multiple signaling pathways, including Smad-dependent signaling and PI3K/PTEN-mTOR signaling. In contrast, another study found that intrathecal delivery of BMP-2 in a spinal nerve ligation rat model did not promote a neuroprotective effect; however, the authors suggested that higher BMP-2 concentrations may be necessary to achieve a therapeutic effect.^45^ Although BMP-2 did not exhibit positive effects in the aforementioned study, BMP-4 was observed to induce allodynia and glial activation. Beyond BMP-2, other BMP family members (BMP-4, BMP-6, and BMP-7) have demonstrated neuroprotective effects both *in vitro* and *in vivo*.^45–48^ Ren *et al.*^49^ explored intracisternal delivery of BMP-7 for stroke treatment in rats and observed significantly enhanced recovery of forelimb and hindlimb placing at early timepoints of delivery. The dual application of BMP’s for both bone regeneration and neuroprotection may address limitations of clinically available products for single-stage TBI treatment. Future studies will focus on evaluating dual BMP-2 delivery strategies to enhance cranial bone regeneration while simultaneously leveraging its neuroprotective potential.

## Conclusion

The current study evaluated the use of a photocrosslinking PHA-TDVT hydrogel with PLGA microspheres containing BMP-2 for the treatment of TBI in a rat *in vivo* model. Covalent bonding between the particles and the PHA polymer, culminating in an interconnected hydrogel, offers an attractive biomaterial for TBI treatment, whereby the biomaterial may accommodate brain swelling while simultaneously supporting cranial bone regeneration. Clinically available regenerative products do not crosslink after placement, which may limit their suitability for TBI treatment due to potential material migration during brain swelling. In contrast, our approach introduces a flexible hydrogel construct that offers a novel treatment paradigm for TBI, with the potential to enable an unprecedented single stage therapeutic strategy. The PHA-TDVT 15% group exhibited desirable mechanical and *in vitro* performance, as demonstrated by yield stress, recovery of G’, compressive modulus, and osteogenic differentiation of hBMSCs. However, overall cranial bone regeneration was limited across all experimental groups, including the PHA-TDVT hydrogel incorporating BMP-2 loaded microspheres. The incorporation of controlled BMP-2 release was initially intended to enhance cranial bone regeneration, but unexpectedly, we observed a neuroprotective benefit in TBI recovery. The PHA-TDVT+µ100 group significantly reduced average motor skill impairment and brain atrophy, highlighting the potential of BMP-2 as a neuroprotective for TBI treatment. Future investigations may be warranted to evaluate dual BMP-2 delivery strategies aimed at simultaneously enhancing cranial bone regeneration and mitigating brain atrophy, advancing the feasibility of a single-stage TBI treatment approach.

## Acknowledgments

The authors would like to thank Dr. Hong Liu and Dr. Yuhua Li at the University of Oklahoma for conducting microcomputed tomography imaging of the rat skulls. Research reported in this publication was supported in part by an Institutional Development Award (IDeA) from the National Institute of General Medical Sciences, the National Institutes of Health, under grant number P20 GM135009. The authors would like to thank Dr. Kar-Ming Fung at the Stephenson Cancer Tissue Pathology Core at the University of Oklahoma Health Sciences Center for providing bone histology services. The authors would like to thank Dr. Mac Harris at the Experimental Pathology Facility at Colorado State University (RRID:SCR_023562) for providing brain histology services. Research reported in the current publication was supported by the National Institute of Neurological Disorders and Stroke of the National Institutes of Health under award number R01 NS123051 and diversity supplement award to Jasmine Deng (R01 NS123051-S1). The content is solely the responsibility of the authors and does not necessarily represent the official views of the National Institutes of Health.

## Author Contribution Statement

<u>Jakob Townsend</u>: Conceptualization, Data curation, Formal analysis, Funding acquisition, Investigation, Methodology, Writing - original draft, Writing - review & editing. <u>Jasmine</u> <u>Deng</u>: Data curation, Formal analysis, Investigation, Methodology, Writing - review & editing. <u>Scott Barbay</u>: Conceptualization, Data curation, Formal analysis, Investigation, Methodology, Project administration, Supervision, Writing - original draft, Writing - review & editing. <u>Brian Andrews</u>: Conceptualization, Funding acquisition, Writing - review & editing. <u>Randolph Nudo</u>: Conceptualization, Formal analysis, Funding acquisition, Investigation, Methodology, Project administration, Resources, Supervision, Writing - review & editing. <u>Michael Detamore</u>: Conceptualization, Formal analysis, Funding acquisition, Investigation, Methodology, Project administration, Resources, Supervision, Writing - review & editing.

